# Multiomics characterization of cell type repertoires for urine liquid biopsies

**DOI:** 10.1101/2023.10.20.563226

**Authors:** Sevahn K. Vorperian, Brian C. DeFelice, Joseph A. Buonomo, Hagop J. Chinchinian, Ira J. Gray, Jia Yan, Kathleen E. Mach, Vinh La, Timothy J. Lee, Joseph C. Liao, Richard Lafayette, Gabriel B. Loeb, Carolyn R. Bertozzi, Stephen R. Quake

## Abstract

Urine is assayed alongside blood in medicine, yet current clinical diagnostic tests utilize only a small fraction of its total biomolecular repertoire, potentially foregoing high-resolution insights into human health and disease. In this work, we characterized the joint landscapes of transcriptomic and metabolomic signals in human urine. We also compared the urine transcriptome to plasma cell-free RNA, identifying a distinct cell type repertoire and enrichment for metabolic signal. Untargeted metabolomic measurements identified a complementary set of pathways to the transcriptomic analysis. Our findings suggest that urine is a promising biofluid yielding prognostic and detailed insights for hard-to-biopsy tissues with low representation in the blood, offering promise for a new generation of liquid biopsies.

## Introduction

Urine is arguably the original liquid biopsy, providing a non-invasive window into the health and functioning of the human body dating back to 4000 BCE^1^. However, current state-of-the-art liquid biopsies are largely focused on nucleic acids in blood plasma. Cell-free RNA (cfRNA) enables a dynamic readout on tissues, organs, and cell types^2,3^ spanning several diseases^4–9^ and physiological changes^9,10^, while cell-free DNA (cfDNA) facilitates noninvasive measurement of growths (i.e. fetus^11,12^ or tumor^13^) and non-self nucleic acids (i.e. microbes^14^ or allograft^15,16^). Urine is collected in the clinic alongside blood; however, the analyte repertoire of this biofluid remains underutilized relative to plasma. Urine is the ultrafiltrate of the blood: some molecules that are present at very low levels in the blood are hyper-concentrated in urine, offering an orthogonal readout to a plasma liquid biopsy. Given that urine interfaces with different tissues from blood and comprises blood waste products, including nucleic acids and metabolites, we hypothesized that measuring the biomolecular repertoire of human urine could facilitate a direct noninvasive window into the health and functioning of cell types and tissues with low representation in blood.

In this work, we characterize the cell type repertoire to form the basis for a urinary liquid biopsy. First, we characterize the landscape of the cell-free and cellular urinary transcriptomes. Second, we compare our findings to the plasma cell-free transcriptome. We then intersect the observable pathways and cell types with untargeted metabolomics data. Altogether, we demonstrate that urine liquid biopsies exhibit a distinct cell type spectrum from blood plasma and identify critical challenges that must be addressed for the adoption of urine liquid biopsy.

## Results

### Characterization of the urine transcriptomic landscape

We first characterized the urine transcriptome (n = 18 patients) (Fig. 1a). Prior studies have focused on RNA in urine sediment^17,18^ or cell-free DNA in urine supernatant^19,20^. Biobanking efforts store various urine fractions^21–24;^ however, a systematic comparison of the information content between the cellular (sediment) and cell-free (supernatant) fractions remains to be made. We therefore isolated and sequenced RNA from both fractions of urine specimens (Methods, Supplementary Fig. 1) from a cohort of patients with kidney stones and healthy control samples from persons without known urinary tract disease (n=12, n = 6 respectively, Supplementary Table 1).

**Fig. 1.**
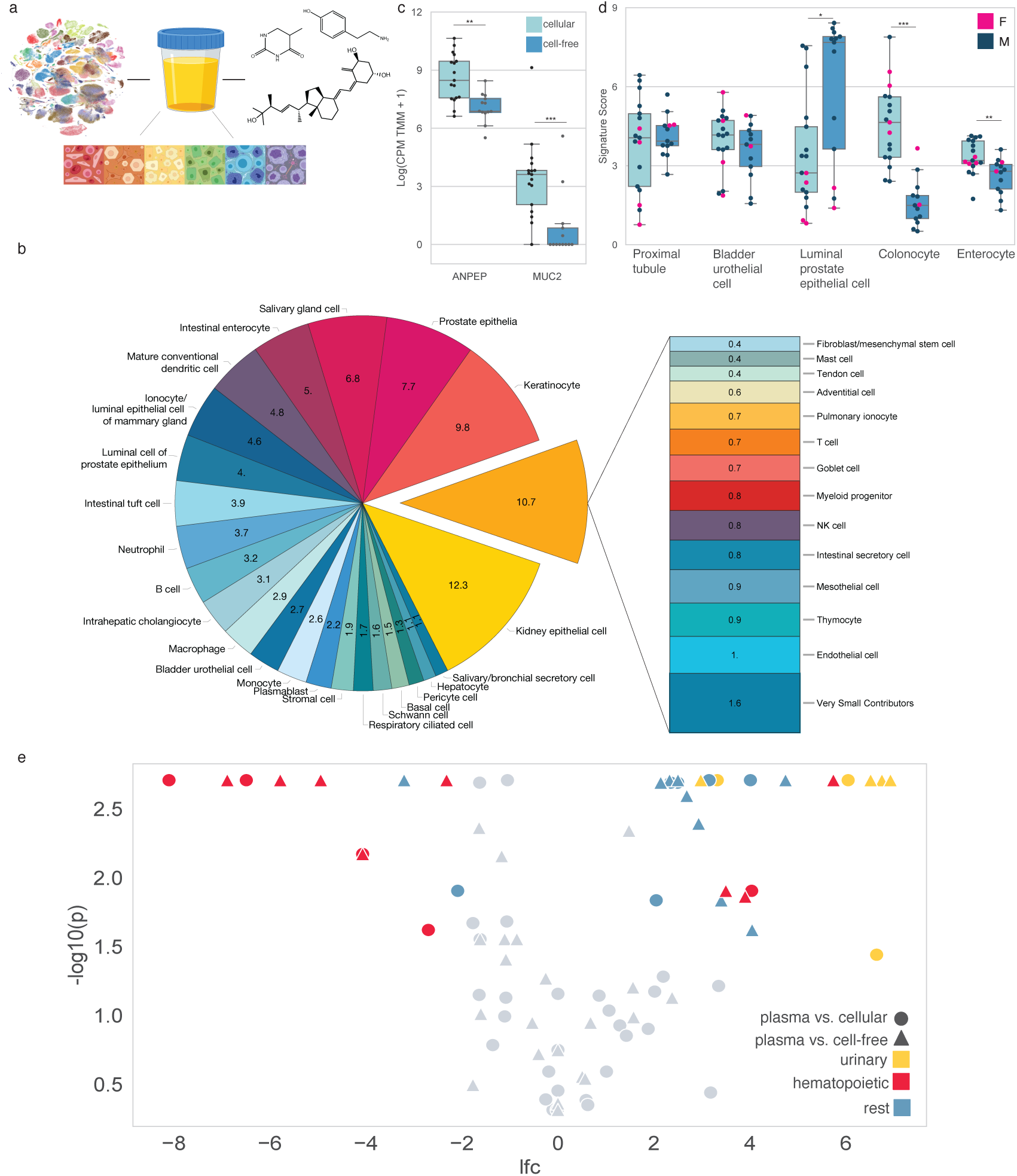
Urine liquid biopsy cell type repertoires are distinct from the plasma cell-free transcriptome. All *P* values were determined by a two-sided Mann–Whitney *U*-test. ** P < 0.01, *** P < 0.001, **** P < 10^-5^. a. Joint transcriptomics and metabolomics to identify the cell types of origin in a urine liquid biopsy b. Mean fractional contributions of cell-type-specific RNA of the urinary cellular transcriptome (n = 4 patients) c. Box plot of intestinal cell type marker gene expression in urine cellular and cell-free RNA (MUC2, p = 6.90 * 10^-4^; ANPEP, p = 1.27 * 10^-3^) d. Signature scores of renal, bladder, prostate, and intestinal cell types in urinary cellular RNA (n = 17 patients) and cfRNA (n = 13 patients) (p = 1.97 * 10^-5^, colonocyte; p = 6.52 * 10^-3^, enterocyte; p = 0.014, luminal prostate epithelial cell) e. Volcano plot of the mean deconvolved fractions of urine cellular RNA (n = 5) or urine cfRNA (n = 5) vs. plasma cfRNA (n = 18 respectively) all from healthy patients.

Deconvolution of these bulk measurements into their underlying relative fractions of cell type specific RNA^3^ using *Tabula Sapiens*^25^, a multi-donor whole-body cell atlas spanning 24 tissues and organs, yielded distinct spectra of cell-type specific RNA (Methods). In urine sediment from healthy controls, we observed predominant fractions from kidney epithelia, prostate epithelia, keratinocytes, and immune cell types (Fig. 1b, Supplementary Figs 2 & 3). Urine directly interfaces with the kidney and prostate, congruent with the observed large relative fractional contributions. Keratinocytes were variable in relative fractional contribution (Supplementary Fig. 2), consistent with the clinical observation that these cells are a source of contamination in conventional urine cytology^26^. Contributions from bladder urothelial cells and Schwann cells were observed, underscoring the ability to measure the whole spectrum of urinary tract cell types. Within a subset of patients with kidney stones, we observed large fractions of neutrophil-specific RNA in their urine cellular transcriptomes, all of whom were dipstick positive for leukocyte esterase (Supplementary Fig. 2c). This observation was consistent with our bulk-level differential expression analysis between the urine cell-free and cellular transcriptomes (Methods): of the urine specimens that were dipstick positive for leukocytes, gene ontology analysis yielded enrichment for immune system processes in cellular urine RNA compared to urine cfRNA (Supplementary Fig 4a). Notably, dipstick urinalysis for leukocytes measures leukocyte esterase, which is released by neutrophils^27^. Neutrophils are one of the first cell types to present in urine during infection, underscoring the sensitivity of the deconvolution method to measure cell type specific signal. In urine cfRNA from healthy male control donors, we observed elevated signal from prostate cell types and secretory cell types (Supplementary Figs. 2 & 3). The prostate comprises several ductal and secretory cells that are responsible for producing seminal fluid and is lined by prostate luminal epithelial cells, which directly interface with prostatic secretions that are the basis for seminal fluid. Urine and seminal fluid pass through the urethra, where it follows that the cell-free fraction of urine from men would comprise dominant contributions of prostate-cell type specific RNA, particularly from the lumen. We additionally observed small relative fractional contributions of cell type specific RNA from non-urinary tract cell types throughout the body, including intrahepatic cholangiocytes, endothelial cells, pericytes, hepatocytes, pulmonary pneumonocytes, and intestinal cell types. In the cfRNA from urinary stone patients, we observed recovery of more cell type-specific signal from non-immune cell types despite being dipstick positive for leukocytes, possibly suggesting that urine cfRNA is more robust to dominant signal associated with inflammation to noninvasively measure urinary tract cell types and tissues.

We were struck by the relative fractional contributions of intestinal cell type-specific RNA across both the urine cfRNA and cellular RNA. We therefore investigated the expression of two intestinal cell type marker genes, *ANPEP* and *MUC2*^28^. *ANPEP* encodes alanyl aminopeptidase, which facilitates peptide breakdown in the small intestine and exhibits highest expression in the intestine across all tissues in the human body. While *ANPEP* is also expressed in renal villi, its expression is highest in the small intestine. *MUC2* encodes a secretory mucin protein that is a primary constituent of gastrointestinal mucus and exhibits expression unique to the intestine^29^. We observed elevated expression of both marker genes in urine cellular RNA relative to urine cfRNA (Fig. 1c), recapitulating the relative abundances of these cell types in our deconvolution results between urine sediment and supernatant.

We then independently assessed the relative abundance of cell type specific signal of renal, bladder, prostate, and intestinal cell types (Fig. 1d, Methods). We observed elevated colonocyte and enterocyte signals in urine cellular RNA relative to cfRNA and elevated prostate epithelia signal relative to urine cellular RNA. We observed no difference in the bladder urothelial and the proximal tubule signatures, suggesting that either urine fraction is conducive to monitoring the cell types in these tissues. Intersection of differentially upregulated genes in urine cellular RNA relative to cfRNA with the cell type-specific gene profiles yielded significant enrichment for luminal prostate epithelial cells (p = 1.43 * 10^-7^, hypergeometric test). Taken together, these abundances are congruent with the original deconvolution results and indicate the presence of intestinal cell types in urine. Altogether, the urinary cellular and cell-free transcriptomes exhibit distinctive landscapes of cell type-specific signal, forming the basis for a noninvasive assay.

### Urine liquid biopsies exhibit distinct cell types of origin from the plasma cell-free RNA transcriptome

Urinalysis is sometimes performed in conjunction with a blood draw. This motivated us to compare the cell type spectra of the urine cellular and cell-free transcriptomes to the plasma cell-free transcriptome (Fig 1e, Supplementary Fig. 4b-f, Methods). We observed that the predominant relative fraction of cell type specific RNA in cellular urine RNA and urine cfRNA originate from cell types of the kidney, bladder, and the prostate. In plasma cfRNA, there is an enrichment for hematopoietic cell types relative to either urinary transcriptome. Independently, we intersected the differentially expressed genes in either urine fraction relative to blood plasma, again observing significant enrichment for bladder urothelial cells, proximal tubules, and prostate luminal epithelia relative to blood plasma (Supplementary Table 2, Methods). Taken together, these findings indicate that the urinary cell type landscape is distinct from blood plasma^3^, underscoring the opportunity to use urine to assay health and disease in different tissues.

Comparison of the pairwise correlations of relative fractional cell type spectra across all urine and plasma samples indicated that both sediment and supernatant samples exhibited higher sample-to-sample heterogeneity than that of plasma (Supplementary Fig. 2c), underscoring an important challenge of the variability in the composition of this biofluid relative to plasma. Differential expression of the urinary cellular and cell-free transcriptomes against the plasma cell-free transcriptome yielded a bimodal distribution in median differentially expressed gene log-fold change (Supplementary Fig 4b, Methods) with a median log-fold change of approximately two in either direction, indicating distinct differences in the compared transcriptional landscapes. Pathway enrichment on genes that were differentially upregulated in either urine fraction relative to plasma (Methods) exhibited striking enrichment for metabolic activity (Supplementary Fig 4c,d), further underscoring the distinct transcriptional landscape of urine from plasma.

### Untargeted metabolomics for a functional readout of the urinary metabolic landscape

Following the observed enrichment for metabolic signals in the urinary transcriptome relative to blood plasma, we sought to examine the metabolite landscape of human urine in conjunction with our RNA measurements. Untargeted LC-MS/MS based metabolomics (Methods) yielded a broad chemical spectrum corresponding to numerous classes of small molecules (Fig. 2a). We observed predominant relative contributions of carboxylic acids and derivatives, organooxygen compounds, benzene and substituted derivatives, and fatty acyls. These findings were broadly congruent with prior work characterizing the human urinary metabolomic landscape^30^ and offered a static slice of the urinary metabolome. While urine metabolomics are frequently performed for biomarker studies^31–33^, a given metabolite can be produced by several different pathways, and metabolomic analyses alone are insufficient for high-resolution insights into underlying pathology.

**Fig. 2.**
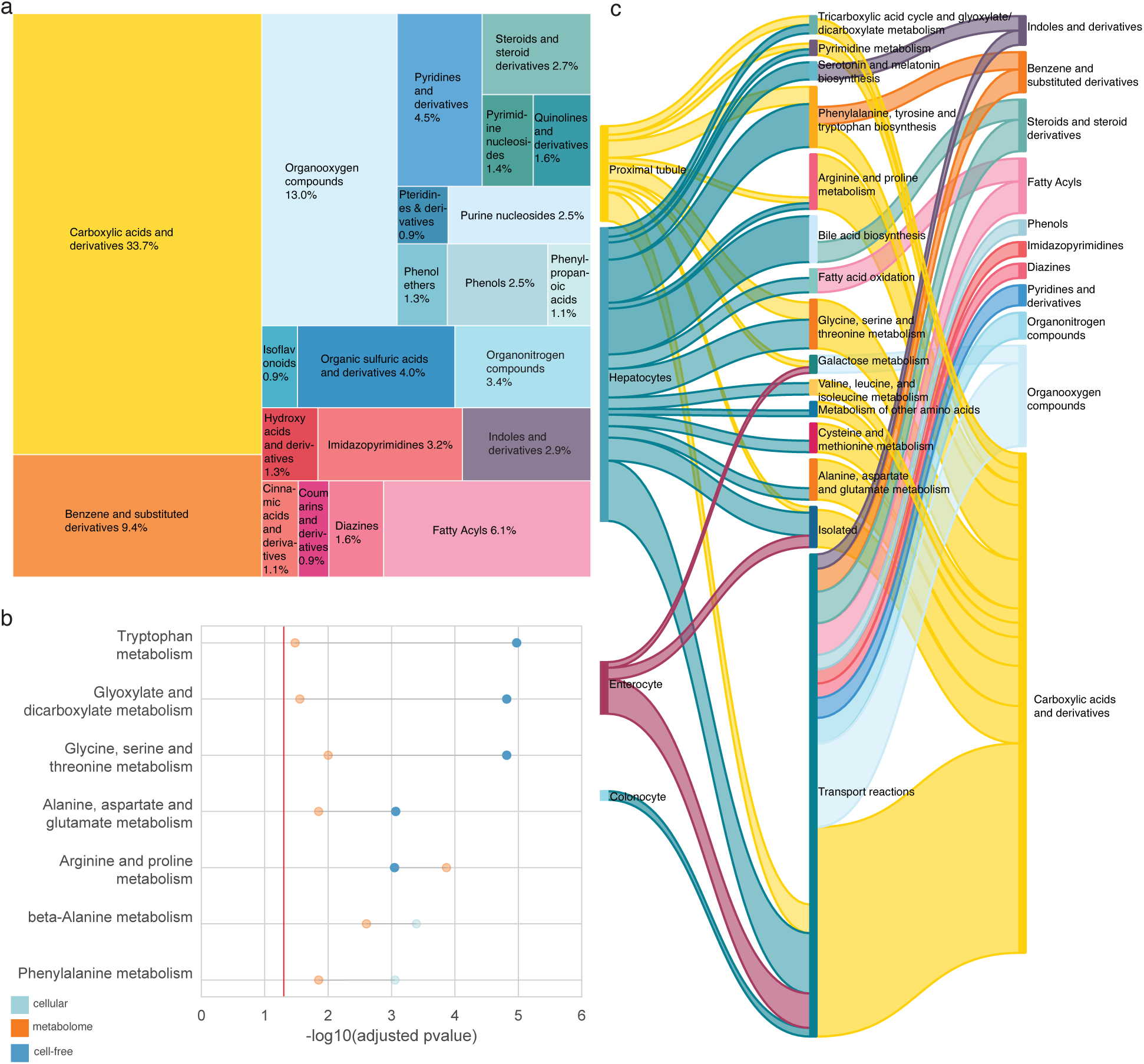
Pathway and cell type level integration of the urine metabolome and transcriptomes. a. Tree map of the chemical classes of molecules measured by HILIC-MS/MS. b. KEGG pathways jointly enriched in urine cellular RNA or urine cfRNA relative to plasma cfRNA and untargeted metabolomics data. c. Sankey plot linking overlapping metabolic pathways from untargeted metabolomics and transcriptomics that were cell type specific. A given metabolic subsystem (middle column) was considered if there were at least two cell type genes mapping to it or five metabolites.

The transcriptome and metabolome are largely studied in isolation; however, the underlying biology is intimately connected *in vivo*^34,35^. While transcriptomics facilitates measurement of changes in gene expression, metabolomics offers a functional readout in the resulting metabolic activity from those gene products. We hypothesized that a joint readout could facilitate better understanding at a pathway level of a disease phenotype, augmenting state-of-the-art liquid biopsy, which focuses on nucleic acids and thereby only offers a limited window into the underlying disease beyond providing a binary sick/healthy readout. Given the rich information within these two ‘omes’, we sought to compare the information content at a pathway level towards applying liquid biopsies as a comprehensive, integrated functional readout on human health.

Pathway enrichment analysis on the measured polar urine metabolites (Methods) included various amino acid metabolic pathways, purine metabolism, and vitamin B6 metabolism (Supplementary Fig. 4g). Comparison of the KEGG-enriched pathways across the transcriptomics and metabolomics data yielded joint enrichment for several amino acid metabolic pathways as well as glyoxylates/dicarboxylates and beta-alanine metabolism, indicating joint measurement of genes and metabolites corresponding to the same pathway (Fig. 2b, Methods).

Given the ability to jointly measure metabolites and transcripts from the same pathway and our observation that various cell types that are enriched in the urinary transcriptome are functionally responsible for metabolic activity, we asked whether overlapping metabolic subsystems were measurable at cell type resolution. To link these measurements of the urinary transcriptome and metabolome, we used the Human Metabolic Atlas (HMA)^36,37^. Cell type gene profile intersections with HMA revealed that the distinct metabolic responsibilities of different cell types were reflected in their cell type specific gene profiles (Fig 2c). For example, the chief responsibilities of the proximal tubule cells are to metabolize amino acids and transport molecules for reabsorption back into the bloodstream, congruent with our observation that proximal tubule cell type specific genes are split between transport reactions and various amino acid metabolic subsystems in HMA. Enterocytes are largely responsible for nutrient absorption into the bloodstream, congruent with the overwhelming proportion of transport reactions for this cell type.

As a principal detoxifier of the human body, hepatocyte-specific genes encode proteins involved in several detoxifying metabolic processes, including drug and xenobiotic metabolism, as well as non-polar molecule metabolism. Intersection of our measured urine metabolome with HMA identified subsets of metabolites corresponding to the same metabolic subsystems as the cell type specific transcriptome measured in urine. Taken together, these findings suggest that joint measurement of the urinary transcriptome and metabolome facilitates an end-to-end readout on gene expression changes and functional responses.

## Discussion

At present, liquid biopsies overwhelmingly focus on blood plasma; however, for organs directly interfacing with other body fluids, the blood may not facilitate sufficient analytical sensitivity on their health and function. Additionally, many diseases of the urinary tract are polygenic^38–40^, oftentimes implicating several pathways for a given pathology; while RNA offers insights into the underlying changes in gene expression, metabolomics provides a functional readout. In this work, we observe that urine liquid biopsies are resolvable at cell-type resolution and are enriched in cell types originating from urological tissues, exhibiting a distinct spectrum of cell type specific signal compared to a blood plasma cfRNA liquid biopsy. We systematically compared the urine cell-free and cellular fractions and observe that urine cfRNA is enriched in prostate cell-type specific signal and is more robust to leukocytes associated with inflammation, suggesting that different urine fractions may be interrogated depending on the disease of interest. The ability to noninvasively resolve bladder urothelial, renal, and prostate epithelia at cell type resolution in urine forms a strong motivation for the development of a much-needed surrogate for the invasive biopsy procedure that incorporates the detailed molecular subtyping information acquired using single-cell transcriptomic atlases^25,41^. We then integrate the metabolome and the transcriptome, demonstrating that these two measurements reflect joint signal from the same pathway.

We hypothesize that our observation of cell type contributions from solid organs outside of the urinary tract (i.e. pneumocytes, enterocytes, cholangiocytes) in urine cfRNA are derived from RNA molecules originating from the blood stream and are filtered by the glomerulus into the urine; notably, the intestine and lung are solid tissues with some of the highest cellular turnovers in the body^42^. Our ability to measure genes from non-urological cell types are congruent with prior reports observing non-urinary tract tumor-derived DNA^43^ as well as for fetal DNA in urine^44^, and underscores the value of deconvolution analyses with a broad cell type space.

We were struck by the elevated prostate cell type-specific signal in urine cfRNA and its potential to address unmet clinical need to identify clinically indolent prostate cancers from those requiring immediate active treatment. We observed that while men exhibited higher prostate-specific gene expression than women, female urine specimens exhibited low, non-zero expression of several nominally prostate-specific genes. Inspection of the distribution of these genes in cell types throughout the whole body with Tabula Sapiens^25^ identified various cell types from other tissues with overlapping functions to the corresponding cell type in the prostate; for example, *SPDEF*, though a known prostate transcription factor, also has known responsibilities in maintaining other mucosal tissues and exhibits expression in goblet cells^45^. *BMPR1B* exhibits comparable median gene expression in the cervix in the Human Protein Atlas RNA consensus dataset^46^ and is expressed in female reproductive tissues^47^. We remark that although these genes exhibit high expression in the prostate relative to other tissues in the body, these genes also exhibit non-zero expression in female tissues and select cell types in other tissues of both sexes.

This study enabled direct comparison of gene expression signal with metabolomic signal in the same sample. Untargeted metabolomics facilitates an unbiased downstream functional readout from gene expression; however, the assignment of measured features to annotated metabolites is database limited; if a molecule is not represented in a database and a known standard does not exist to validate chemical identity, the assertion of its presence is limited^48^. We observed metabolites enriched in the same pathways we observed in the urinary transcriptome, and we further observed that metabolic cell types whose signal we measure with RNA participate in distinct metabolic pathways with measurable metabolites. We are optimistic that the ability to use metabolomics as a functional readout alongside gene expression changes in linking genotype to disease phenotype and enactment will improve with the development of new tools that expand the space of identifiable compounds from untargeted metabolomics data^49,50^ and the continued joint measurement of the transcriptome and metabolome, thereby offering increased potential for more precise biomarker panels for disease diagnostics and subtyping.

This work also identifies and addresses challenges facing the development of urine liquid biopsies. A given specimen may be contaminated during initial collection, sample storage, and/or processing. Our observation of variable keratinocyte signal in urine may underscore the importance of clean catch during patient sample collection. We addressed challenges of solute concentration by using RNA-library normalization that directly accounts for variability in library size that can skew differential comparisons^51^ and measured spot urine creatinine, which is excreted in approximately constant amounts^52^, to control as a covariate in differential expression. The urine composition relative to the plasma cell-free transcriptome underscores the variability in this biofluid (Supplementary Fig. 2); however, given the proximity of urine to hard-to-biopsy tissues with low abundances in the blood and the ability to measure elevated cell type specific signal, we anticipate that urine may be readily used for noninvasive measurement of the health and functioning of urinary tract cell types.

Our study indicates that urine liquid biopsy measurements facilitate direct, non-invasive access to unique cell types and tissues that are challenging to access with conventional needle biopsy and reflect urinary tract tissue-specific pathophysiology that are not readily measurable in a plasma liquid biopsy. We anticipate that urine liquid biopsies and the use of single-cell transcriptomic atlases as a reference to interpret these measurements offer a promising new frontier for the fields of liquid biopsy and noninvasive diagnostics.

## Acknowledgements

This work is supported by the Chan Zuckerberg Biohub. SKV is supported by the NSF Graduate Research Fellowship (Grant # DGE1656518), the Benchmark Stanford Graduate Fellowship, and the Stanford ChEM-H Chemistry Biology Interface (CBI) training program. JAB was supported by a National Institutes of Health Ruth L. Kirschstein NRSA F32 Postdoctoral Fellowship (1F32GM134689).

J.C.L. is supported by NCI R01 CA244526 and Department of Veterans Affairs Merit Review I01 BX004962. We thank C. Liou, N. Mehta, J. Long, E. Sattely, and Y. Dai for metabolomics discussions. We thank J. Zou for confidence interval discussions. We thank B. Yu for sequencing discussions. We thank T. Meyer for renal discussions and urine specimens. We thank Quake lab members past and present for helpful discussions. We thank the ChEM-H Knowledge Center for QTOF access. We thank R. LaMantia for making the rainbow cell spectrum in Fig 1A; the cells in figure 1A were generated using BioRender.

## Author contributions

SKV conceptualized this study. SKV designed the study with input from BCD, JAB, GBL, and SRQ. BCD, JAB, IG performed untargeted metabolomics; BCD performed metabolomic data preprocessing. SKV and HJC developed cellfracker. RY performed sequencing. SKV performed all analyses with metabolomics analysis input from BCD. TJL, VL, KEM, JCL collected clinical specimens. SKV, BCD, JAB, KM, JL, RL, GBL, CRB, SRQ interpreted results. SKV and SRQ wrote the manuscript. All authors revised this manuscript and approved it for publication.

## Data deposition

Raw sequencing data will be made available on SRA. Raw metabolomics data will be made available on Metabolights. All code will be made available on github.

## Code availability

Code for the work in this manuscript will be made available on GitHub. Cellfracker is openly available at github.com/sevahn/cellfracker.

## Methods

### 1. Sample collection and preprocessing

#### 1.1 Urine specimen collection

Clean catch urine specimens were collected from patients with kidney stones (n = 12) and healthy controls without known kidney disease (n = 6). Written informed consent was obtained from all participants. The study was approved by the Stanford Institutional Review Board and was conducted in accordance with the Declaration of Helsinki.

#### 1.2 Urine specimen preparation and processing

Voided specimens were stored at +4°C until processing; samples were processed within 6 hours of collection. Dipstick analysis was performed on a separate aliquot (Fisherbrand, 23-111-262). Whole urine was aliquoted for metabolomic analysis. The remaining sample was spun at 4°C and 3000g for 30 min. Sediment samples were prepared as previously described^53^: 0.1% v/v betamercaptoethanol (Millipore, 444203) and 1 mL Trizol (Ambion, 15596026) were added to the pellet following centrifugation. Samples were frozen at −80 °C for subsequent processing. Spot creatinine was measured using an aliquot of frozen urine (Biotechne, KGE005).

#### 1.3 Creatinine Factors

impacting urine solute concentration include voided volume and patient hydration^52,54^, thereby possibly confounding abundance quantification of a given analyte. This is unlike blood plasma, a homogenous biofluid of approximately uniform solute concentration. As a proxy for evaluating patient hydration, spot creatinine was measured^52^. Standard urine dipstick (Fisherbrand, 23-111-262) was additionally measured. This basic specimen workup then provided a baseline from which we could compare the results of our high-resolution measurements in our study.

### 2. Experimental

#### 2.1 RNA isolation

Urine sediment RNA isolation was performed as previously described^18,39^. Briefly, frozen Trizol pellets were thawed on ice and isolated according to manufacturer’s instructions (Qiagen, 74104). Urine supernatant RNA samples were thawed at room temperature. Cell-free RNA was isolated from between 2-8 mL following manufacturer’s instructions (Qiagen, 55114). Isolated RNA was DNase-treated (Lucigen, DB0715K), cleaned and concentrated (Zymo, R1015), then quantified (Agilent, 5067-1513).

#### 2.2 Library preparation and sequencing

All libraries were prepared using the SMARTer Stranded Total RNaseq Kit v2 – Pico Input Mammalian Components (634419, Takara) according to manufacturer’s protocols and barcoded (634452, Takara). Urine samples were then sequenced with (either the 2 x 150 bp or 2 x 75 bp configuration) to a mean depth of 30 M reads per sample on NextSeq 2000 (Illumina Inc); sequencing length did not impact gene detection efficiency (Supplementary Fig. 1b).

#### 2.3 Metabolomics sample preparation

Urine specimens were thawed on wet ice, 100 µL aliquots were extracted in 1.5 ml Eppendorf polypropylene tubes (022363204, Eppendorf) with addition of 80 uL of chilled extraction solvent containing stable deuterated internal standards (Supplementary Table 3) at − 20°C (1:1 ACN:MEOH with 1% Water) followed by an additional 320 uL of 1:1 ACN:MEOH at −20°C. Specimens were hand shaken to mix, then chilled at −20°C for one hour. Next, specimens were vortexed for 10 seconds and centrifuged at −9°C for 5 minutes at 14000 RCF. The supernatant was then transferred to a fresh tube for drying in a centrivap at room temperature. Residues were then reconstituted in 100uL of 3:2 ACN:H2O containing 60ng/mL CUDA (1-cyclohexyl-urido-3-dodecanoic acid). Specimens were then vortexed, centrifuged for 10 seconds at 14,000 RCF, from which the supernatant was sealed in glass autosampler inserts (C4011631, Thermo Scientific), and promptly injected onto a Waters Acquity UPLC BEH Amide column (150 mm length × 2.1 mm id; 1.7 μm particle size) with an additional Waters Acquity VanGuard BEH Amide pre-column (5 mm × 2.1 mm id; 1.7 μm particle size) maintained at 45°C and coupled to an Thermo Vanquish UPLC. Mobile phases were prepared with 10 mM ammonium formate and 0.125% formic acid and 100% LC-MS grade water for mobile phase (A) or (B) 95:5 v/v acetonitrile:water. Gradient elution:100% (B) at 0–2 min to 70% (B) at 7.7 min, 40% (B) at 9.5 min, 30% (B) at 10.25 min, 100% (B) at 12.75 min, isocratic until 16.75 min with a column flow of 0.4 mL/min.

#### 2.4 Metabolomics instrumentation

Metabolites were measured using a hydrophilic interaction liquid chromatography (HILIC) column coupled to a Thermo Q Exactive HF Hybrid Quadrupole-Orbitrap mass spectrometer. Full scan MS1 and data dependent MS2 (MS-ddMS2) were collected in both positive and negative mode ionization (separate injections). MS^2^ selection was prioritized for metabolites with known mass to charge ratios and, LC-MS/MS method specific, retention times using an inclusion list from an in-house library generated from authentic standards. Additional MS^2^ data were collected in a data-dependent manner when scan bandwidth was available. Mass range was 60-900 *m/z*. MS^1^ resolution was 60k; MS^2^,15k. Data was collected in centroid mode, loop count was 4, quadrupole isolation window was 1.0 Da.

### 3. Data Preprocessing

#### 3.1 RNA

Reads were trimmed (trimmomatic version 0.36) and mapped (STAR version 2.7.3a) to the human reference genome (hg38). Duplicate reads were then marked and removed (MarkDuplicates version 4.1.1). Mapped reads were then counted (htseq-count version 0.11.1). Read statistics were estimated using FastQC (version 0.11.8). The snakemake (version 5.8.1) workflow management system was used to perform the above and MultiQC (version 1.7) was used to aggregate performance statistics. 3’ bias, intron to exon ratio, and ribosomal read fraction and were computed. The associated sample-level QC cutoffs were then applied as previously described to the rounded metrics^55,56^. The cutoff ribosomal fraction was set to 0.5 given that samples were sequenced to a depth that a sufficient number of genes were detected and these samples were not PCA outliers.

#### 3.2 Metabolomics

Metabolomics data were processed using MS-Dial (version 4.60)^57^ for peak picking, alignment, and annotation. Metabolites were annotated by accurate mass and MS^2^ spectral matching. Additionally, when a metabolite’s method specific retention time was known, retention time matching +/- 0.4 minutes was required. MS^2^ reference spectra were obtained from MassBank of North America (https://mona.fiehnlab.ucdavis.edu/)^58^ or in-house analysis of authentic standards^59^. MS1 accurate mass tolerance was 0.01 Da, MS2 tolerance was 0.015 Da. All annotations were manually reviewed.

### 4. Cell type deconvolution

#### 4.1 Cell type deconvolution method

Samples were deconvolved using nu support vector regression as previously described^3^. Briefly, given some reference of the gene expression profiles of the cell types in Tabula Sapiens (e.g. basis matrix, *A*), the objective is to learn the relative fractions of the cell types to recapitulate the signal in the bulk mixture *b*:

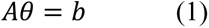

Resulting fractions of cell type specific RNA (*θ*) correspond to the normalized positive coefficients learned by the best model (defined as the model yielding the smallest RMSE) across a grid search of nu and C values.

To facilitate the transparent deconvolution of a user’s samples and interpretation of deconvolution results, we created an open-source, command line tool called cellfracker (cell fraction estimator). A given run outputs the cell type fractions as well as the support vectors used, facilitating transparent interpretation of deconvolution results. To deconvolve your own samples with Tabula Sapiens or a prespecified user’s basis matrix, a pip installable version is available online or at github.com/sevahn/cellfracker.

#### 4.2 Bootstrap procedure

We bootstrapped 90% confidence intervals for a given cell type fraction in a given sample by randomly sampling genes and their associated counts with replacement from a given sample^57^ until total read count was within 10% of the original. Deconvolution was performed described in the section entitled ‘Cell type deconvolution method’.

Next, the 90% bootstrapped confidence interval for a given cell type fraction in a given sample was determined:

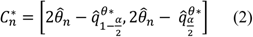

Where *n* corresponds to the number of samples; 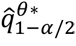 and 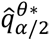 correspond to the *α*/2 and 1 - *α*/2 quantiles from given confidence level *α*, and 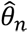 is the estimated valued from the collected data.

#### 4.4 Comparison of deconvolved cell type fractions of the plasma and urinary cell-free transcriptomes

Deconvolved relative fractions of cell type specific RNA in either urine fraction were compared to deconvolved plasma samples from a previously published study^11^ (non-cognitively impaired cohort, BioIVT sample source) with a two-sided Mann Whitney U test. As with the differential expression analyses, a given cell type were only compared if the proportion of nonzero coefficients was greater than or equal to the smallest relative proportion of samples of a given type. Multiple hypothesis testing correction was performed using the Benjamini Hochberg procedure with a minimum significance cutoff level of 0.05.

#### 4.5 Cell type gene profiles for signature scoring

Cell type gene profiles in context of the whole body for were derived from datasets for the intestine^28^, prostate^41^, and bladder^25^ as previously described^9^. Remaining cell type gene profiles were as previously reported^9^. Scanpy^58^ (version 1.8.1) was used to perform the single cell differential expression and the NX values from the Human Protein Atlas RNA consensus dataset^44^ were used to determine gene expression specificity in context of the whole body using the Gini coefficient^59^.

#### 4.6 Signature scoring

The signature score for the j^th^ sample is given by:

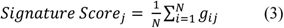

Where *N* is the number of genes in the cell type gene profile and *g*_+)_ is the log-transformed, counts per million and TMM normalized gene expression values. Counts tables used for signature scoring were filtered as described in section 5.1.

### 5. RNA-seq differential expression

Two sets of comparisons were performed: (1) urine cellular RNA vs urine cfRNA and (2) urine RNA vs. plasma RNA. For the following subsections, the described analyses were repeated separately for each of these two comparisons.

#### 5.1 Gene filtering and normalization

Genes used in all comparative analysis following deconvolution were expressed above 1 log-normalized CPM counts in at least the proportion of samples equal to the minimum group sample size^60^. Unlike urine cfRNA, urine cellular RNA originates from cells and is intact and must be fragmented during library preparation. While samples were sequenced together and to the same approximate depth, a given gene in a sediment sample may have more unique raw counts than the same gene in a corresponding urine supernatant sample because of the fragmentation library preparation step. To this end, samples and the corresponding filtered genes were then TMM normalized^51^ to account for variations in library size. The resulting median gene expression of a list of housekeeping genes^61^ were then compared; no significant differences were observed in the median housekeeping gene value (p = 0.447, cellular urine RNA vs. plasma cfRNA; 0.111, urine cfRNA vs. plasma cfRNA; 0.447, cellular urine RNA v. urine cfRNA; two-sample t-test with Benjamini-Hochberg correction) (Supplementary Fig. 1f).

#### 5.2 Differential expression

Differential expression on the RNA-seq data^60^ was performed using limma (version 3.52.2)^61^ and edgeR (version 3.38.4)^62^. Two sets of differential expression were performed: (1) to compare the urine cell-free and cellular transcriptomes and (2) to compare either urine transcriptome to the plasma cell-free transcriptome. In both comparisons, a means model was fit to the data. In the first comparison, the explanatory variables were urine fraction (cell-free or cellular) and the leukocyte dipstick status (binarized positive or negative for leukocytes on dipstick urinalysis). These two factors were converted to a single factor. Volume and spot creatinine were treated as covariates; patient sex and whether or not the sample came from a healthy control were treated as additional factors. In the second comparison, the two explanatory variables were biofluid (cell-free urine, cellular urine, cell-free plasma) and whether or not the sample came from a healthy control. These two factors were converted to a single factor. Sample volume was treated as a covariate. Leukocyte dipstick status and patient sex were treated as additional factors.

The function ‘voomWithQualityWeights’^63^ was run with “genebygene” estimation. Across both sets of differential expression analyses, samples originating from the same patient were controlled for as a random effect using the ‘duplicateCorrelation’^64^ function to estimate the inter-subject correlation. For each of these two differential expression analyses, since multiple contrasts were simultaneously tested, to address multiple hypothesis testing across all considered genes, the function ‘decideTests’ with the ‘global’ method was applied to identify the differentially expressed genes for a given contrast. Multiple hypothesis testing correction was performed using the Benjamini Hochberg procedure with a minimum log2 fold change of 0 and a minimum significance cutoff level of 0.05.

#### 5.3 Bootstrapping confidence intervals

Of the differentially expressed genes passing the multiple hypothesis correction, 95% confidence intervals were bootstrapped to estimate the log fold change by sampling with replacement and computing the median expression value (equation 3). The data were log-transformed and normalized as per the limma ‘cpm(log = TRUE)’ function. A total of 1000 bootstrap replicates were computed per differentially expressed gene for a given contrast. A gene whose bootstrapped confidence interval spanned positive and negative values was discarded. Stated differently, a gene was kept as differentially expressed if and only if the bootstrapped confidence interval spanned positive values if the true log fold change was greater than zero or the bootstrapped confidence interval spanned negative values if the true log fold change was less than zero.

### 6. Pathway and cell type enrichment analysis

Pathway enrichment analyses were restricted to the Kyoto Encyclopedia of Genes and Genomes (KEGG) for comparison of the same pathway between the transcriptome and metabolome. For the transcriptomic data, pathway enrichment analysis was performed using g:Profiler^65^; metabolomics data, metaboanalyst 5.0 ^66^. As g:Profiler returned adjusted p values, uncorrected p values were computed using a hypergeometric test on the returned effective domain size, query size, intersection size, and term size. Multiple hypothesis correction then was performed across all raw p-values using a Benjamini Hochberg test with alpha = 0.05 (statsmodels version 0.10.1).

For transcriptomic and metabolomic pathway enrichment, the respective sets of all genes and metabolites in KEGG were used as the background. We note that the background used in untargeted metabolomics can strongly influence the resulting pathway enrichment results^67^. This contrasts with a transcriptomic dataset, where samples sequenced to sufficient depth will exhibit comprehensive coverage of the transcriptome; a high-confidence metabolites identified with untargeted metabolomics covers a comparatively smaller fraction of the total metabolome. However, we note this limitation and assert that the basis for this methodological choice in background are the high confidence in the detected metabolite identities resulting from the precision metabolomics experiment, the multiple hypothesis correction strategy, and that the enriched pathways are congruent with independent studies of the normal urine metabolome^30^, thereby limiting the detection of spurious enriched pathways in the urinary metabolome.

Cell type enrichment analyses were performed using the space of protein coding genes as the background, differentially upregulated genes in the urine cellular or cell-free fraction vs. plasma as the query, cell type specific gene profiles as the reference. Multiple hypothesis correction was performed as done for the pathway enrichment across p-values for both urine fractions.

### 7. Metabolomics

#### 7.1 Chemical compound mapping

Chemical taxonomy on the identified metabolite InChiKeys were determined using ClassyFire^68^ (version 1.0). Visualization was performed using squarify (version 0.4.3).

#### 7.2 Integrated cell type and metabolite mapping

The human metabolic atlas^36,37^ (version 3.1) was used to map the metabolite reactants and products or a given gene to a given reaction and subsequent pathway. Given the number of synonyms for a given metabolite, metabolites in the human metabolic atlas were mapped to InChiKeys programmatically using pubchempy (version 1.0.4). The InChiKeys of these metabolites were intersected with those of the metabolites detected; the first 14 characters^69^ were considered given that this encodes the molecular constitution of a compound. Ensembl gene ID were intersected between cell type specific gene profiles (defined in section 4.5) and Ensembl gene ID of genes in HMA. A given metabolic subsystem (middle column) was considered if there were at least two cell type genes mapping to it or five metabolites. Sankey plot visualization was performed using holoviews (version 1.14.9).

---

**Supplementary Fig. 1.**
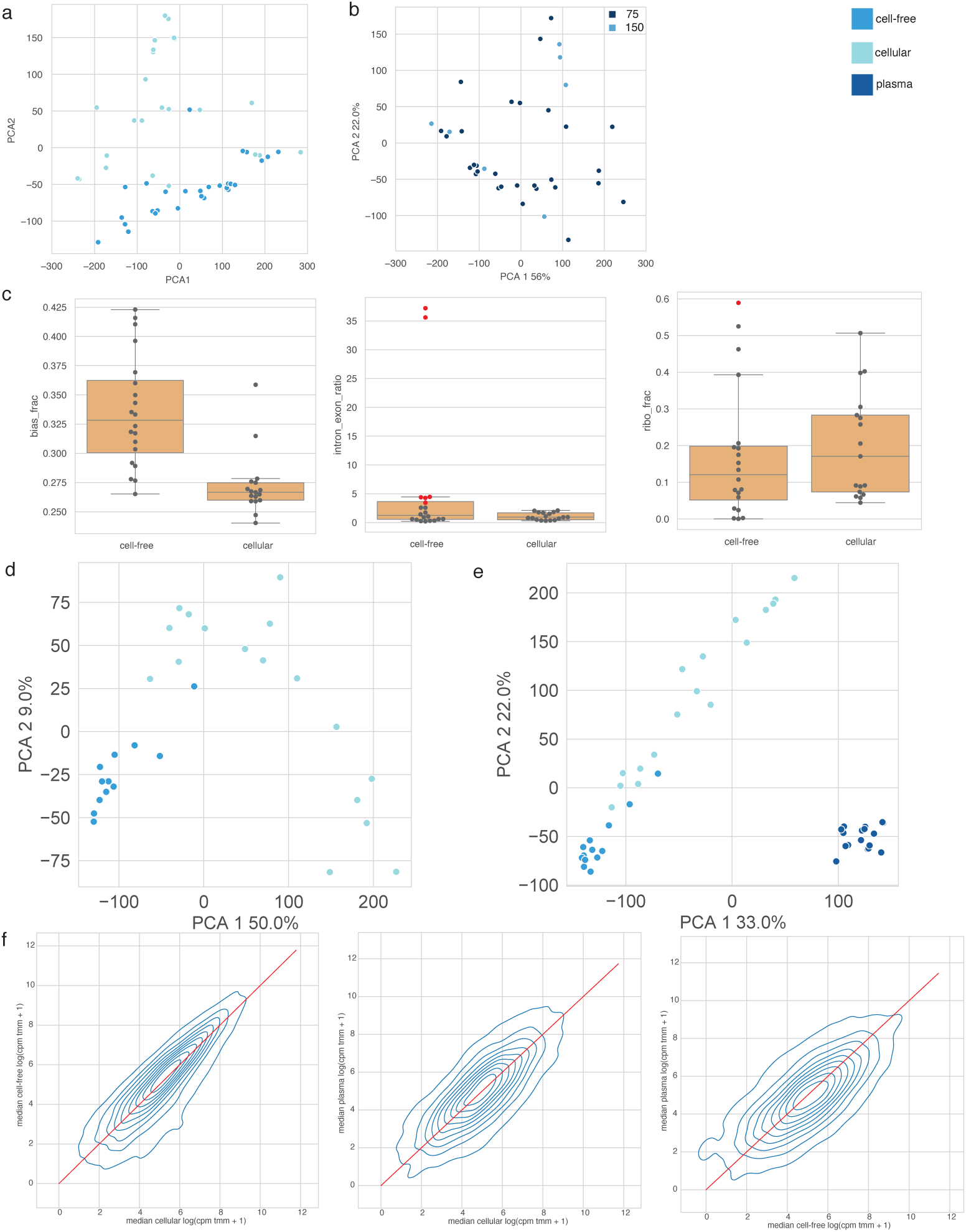
Transcriptomics sample quality control. a. PCA of sequencing data prior to QC filtering b. PCA of sequencing data colored by read length c. Quality control metrics (3′ bias fraction, ribosomal fraction, and DNA contamination) were determined for each sample. Samples with outlier values are highlighted in red and were not considered in subsequent analyses (see Methods section 3.1 “RNA data preprocessing”) d. PCA plot of urine cellular and cell-free fractions normalized counts following sample filtering and normalizing e. PCA plot of urine cellular, urine cell-free, and plasma normalized counts following sample filtering and normalizing f. Pairwise 2D density plots of median housekeeping gene value for a given biofluid pair. Red line demarcates the line y = x.

**Supplementary Fig. 2:**
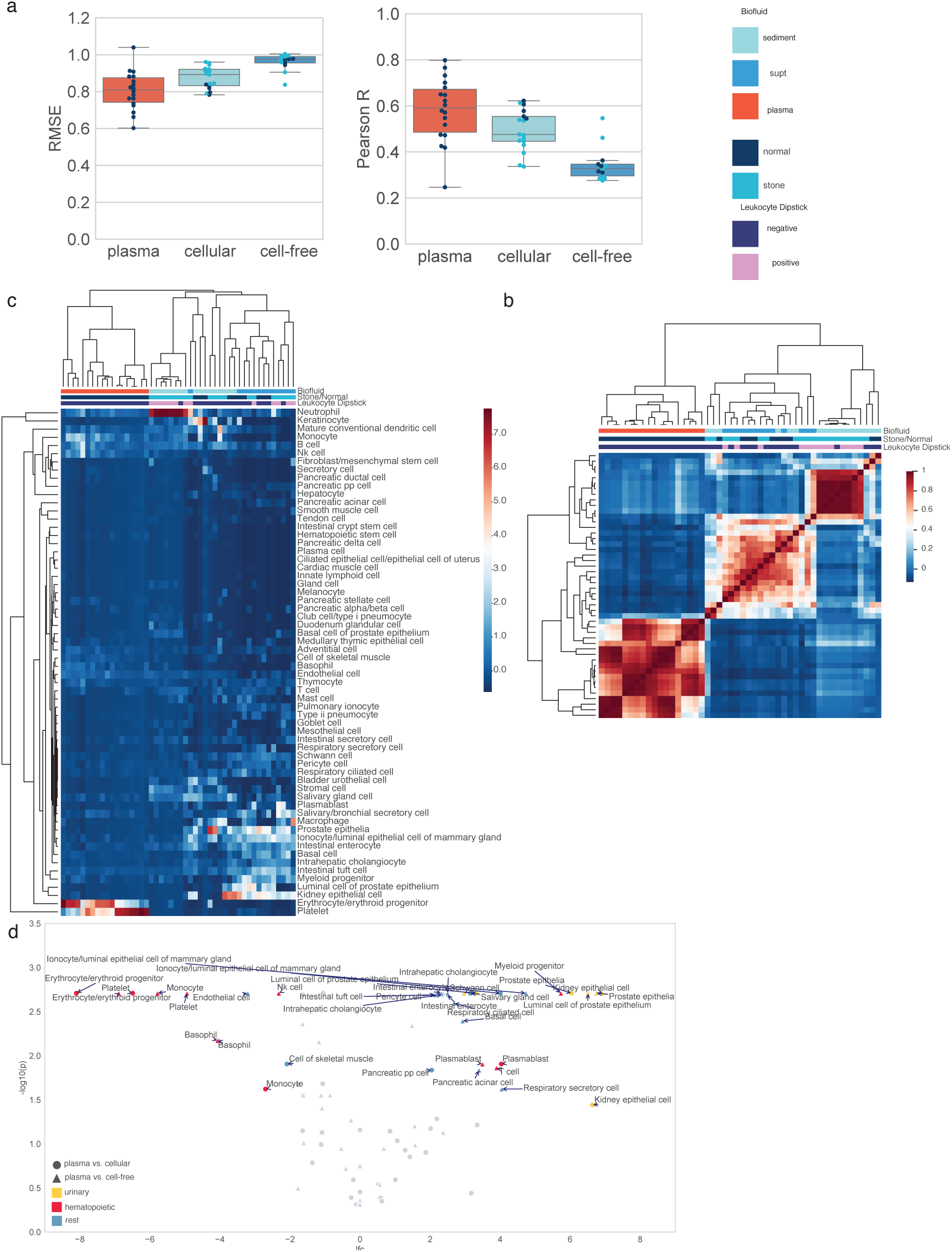
nuSVR decomposition of the urine cell-free and cellular transcriptomes with Tabula Sapiens. a. Deconvolution RMSE and Pearson correlation between predicted vs. measured expression for all samples. b. Complete linkage cluster map of pairwise Pearson correlation of deconvolved cell type fractions across all samples (n = 48) c. Complete linkage cluster map of relative deconvolved fractions of cell-type specific RNA across all samples (n = 48) d. Annotated volcano plot from Fig 1e; see Methods section 4.4 “Comparison of deconvolved cell type fractions of the plasma and urinary cell-free transcriptomes”

**Supplementary Fig. 3.**
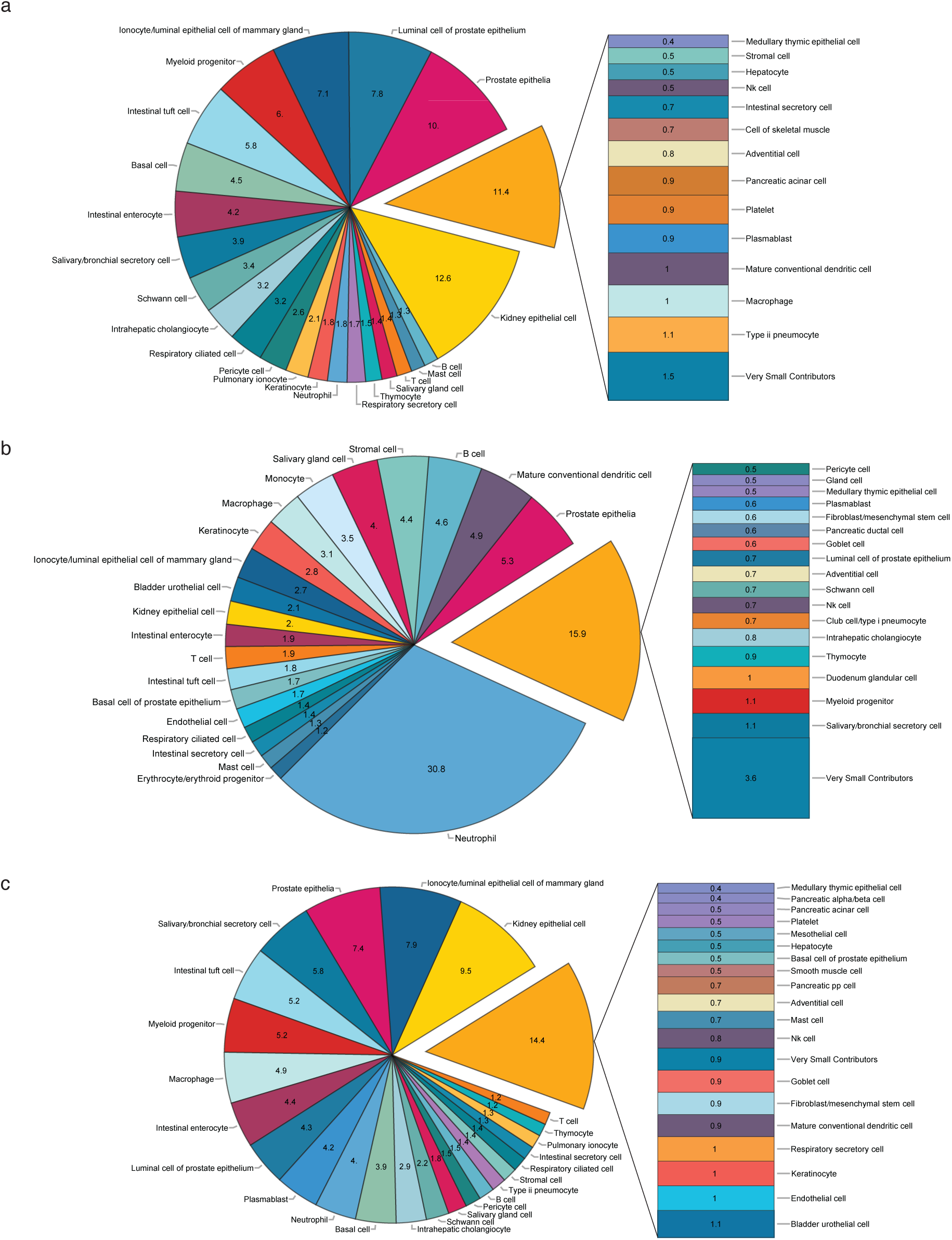
Mean deconvolution pie charts of urine cell-free and cellular samples. a. Urine cfRNA from normal patients (n = 5) b. Urine cellular RNA from stone patients (n = 12) c. Urine cfRNA from stone patients (n = 8)

**Supplementary Fig. 4.**
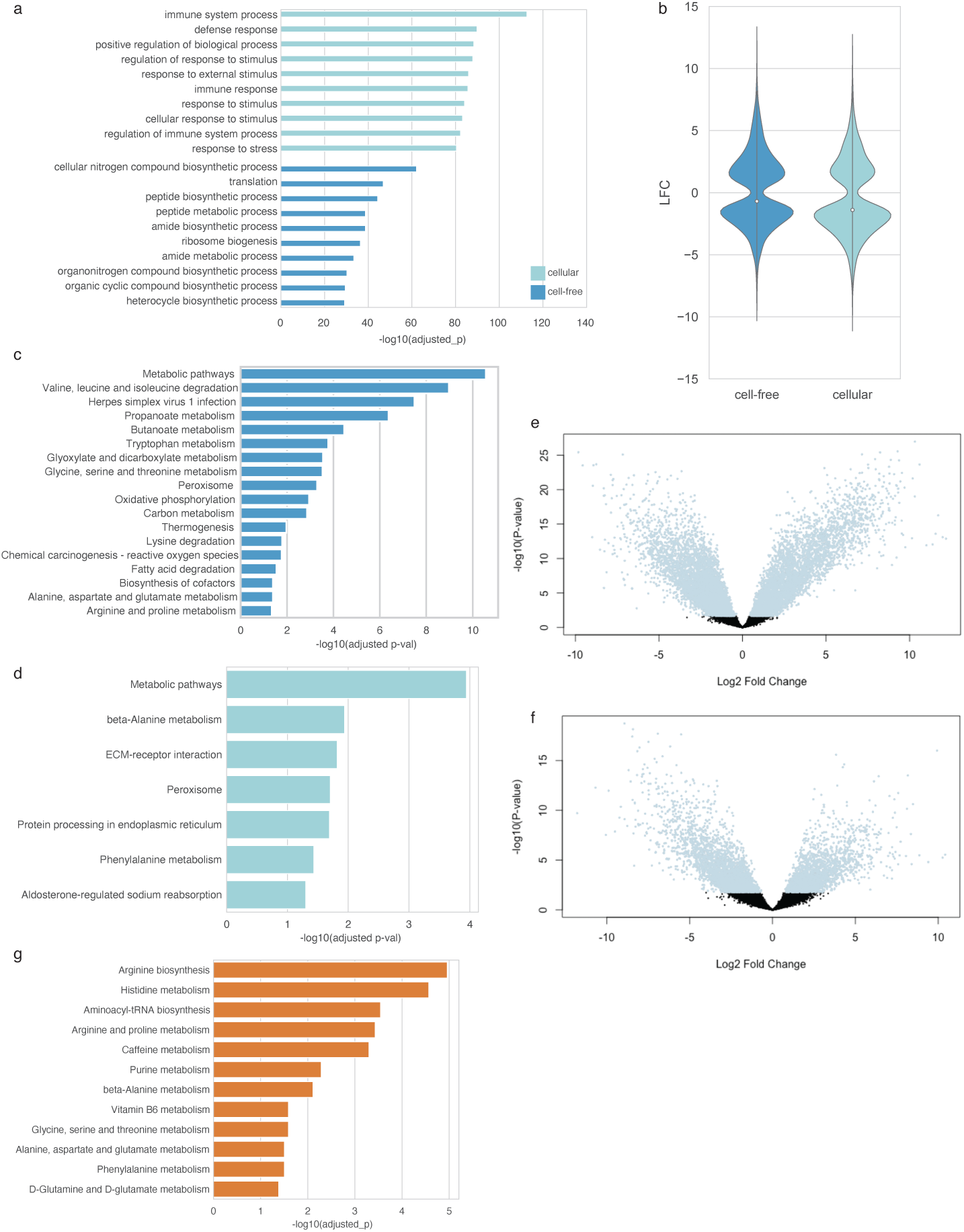
Bulk-level differential expression of urine cellular, urine cell-free, and plasma cell-free transcriptomes. a. Pathway enrichment (Gene Ontology: Biological Process) on cellular vs. cell-free fractions of kidney stone patients b. Violin plot of log2 fold change of differentially expressed genes between the plasma cell-free transcriptome and the respective urine transcriptomes c. KEGG pathway enrichment on genes upregulated in the healthy urine cell-free transcriptome relative to the plasma cell-free transcriptome d. KEGG pathway enrichment on genes upregulated in the healthy urine cellular transcriptome relative to the plasma cell-free transcriptome e. Volcano plot of differential expression contrast in 2c. Positive LFC corresponds to genes upregulated in plasma; negative, genes upregulated in urine cfRNA. f. Volcano plot of differential expression contrast in 2d. Positive LFC corresponds to genes upregulated in plasma; negative, genes upregulated in cellular urine RNA. g. KEGG pathway enrichment on pathways enriched in measured metabolites.

**Supplementary Fig. 5.**
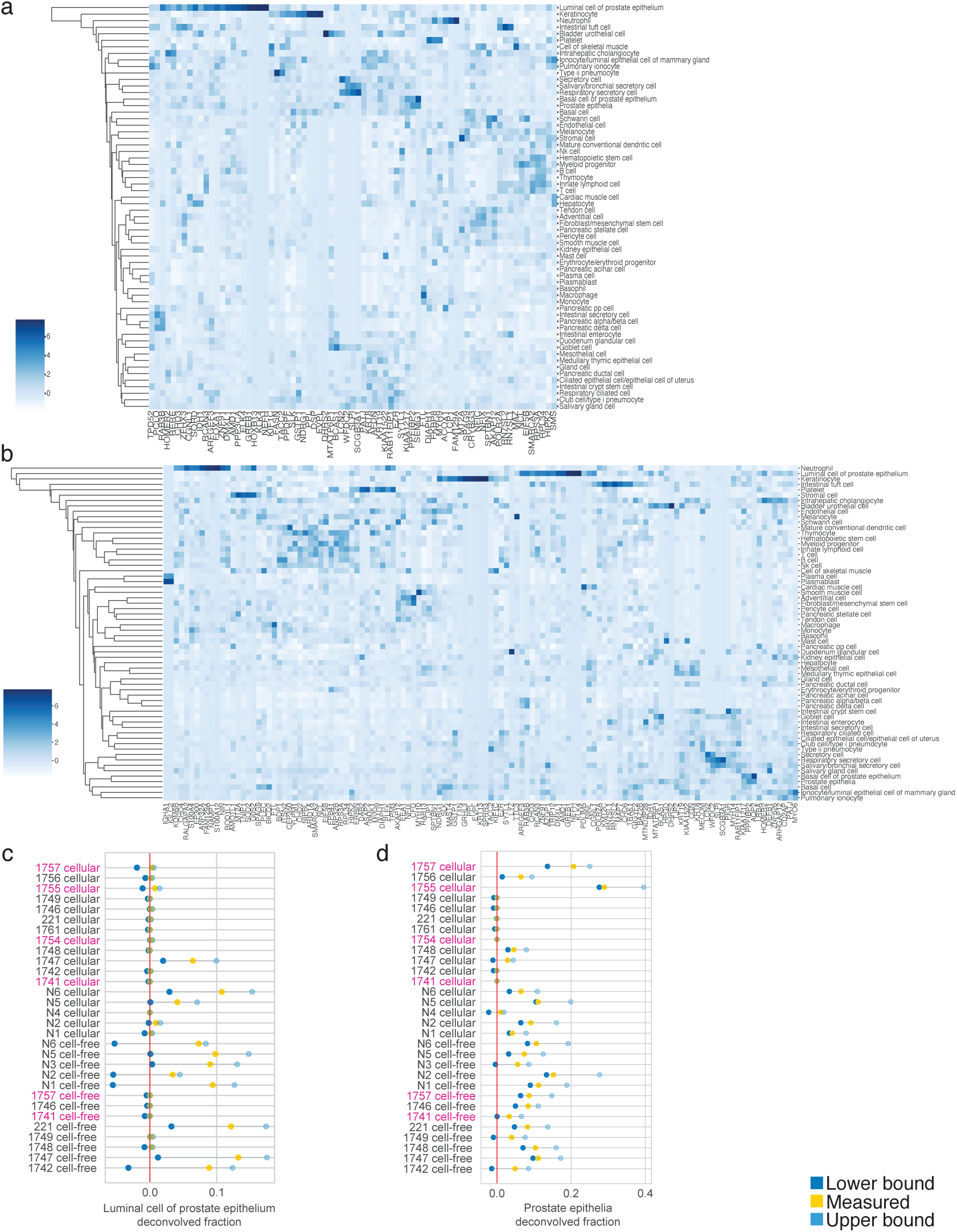
Visualization of prostatic gene expression. a. Linkage clustermap of relative (Z-scored) gene expression in Tabula Sapiens basis matrix V1 for genes with predicted expression greater than 2 standard deviations across normal urine cfRNA samples (n = 5 patients). b. Linkage clustermap of relative (Z-scored) gene expression in Tabula Sapiens basis matrix V1 for genes with predicted expression greater than 2 standard deviations across across ureter stone urine cfRNA samples (n = 8 patients). c. Bootstrapped 90% confidence interval on deconvolved relative fractions of prostate epithelia and luminal cell of prostate epithelia cell type specific RNA (Methods)

## Supplementary Information for Vorperian et al ‘Multiomics characterization of cell type repertoires for urine liquid biopsies’

**Supplementary Table 1:** Summary of patient information of specimens passing QC.

| Patient ID | Sex | Age | Condition |
| --- | --- | --- | --- |
| 1741 | F | 35 | Kidney stones |
| 1742 | M | 30 | Kidney Stones |
| 221 | M | 73 | History of kidney stones |
| 1746 | M | 73 | Kidney stones |
| 1747 | M | 60 | History of kidney stones |
| 1748 | M | 79 | History of kidney stones |
| 1749 | M | 77 | History of kidney stones |
| 1754 | F | 48 | Kidney stones |
| 1755 | F | 82 | Kidney stones |
| 1756 | M | 74 | Bladder inflammation |
| 1757 | F | 74 | Kidney stones |
| 1761 | M | 77 | Bladder stone |
| N1 | M | N/A | Healthy control without known urinary tract disease |
| N2 | M | N/A | Healthy control without known urinary tract disease |
| N3 | M | N/A | Healthy control without known urinary tract disease |
| N4 | M | N/A | Healthy control without known urinary tract disease |
| N5 | M | N/A | Healthy control without known urinary tract disease |
| N6 | M | N/A | Healthy control without known urinary tract disease |

**Supplementary Table 2:** Enrichment of cell types in differentially upregulated urine genes relative to the plasma cell-free transcriptome.

| Cell type | Biofluid | Adjusted p |
| --- | --- | --- |
| bladder urothelial cell | cellular urine RNA vs. plasma cfRNA | 3.70 E-05 |
| Proximal tubule | cellular urine RNA vs. plasma cfRNA | 1.52 E-09 |
| Luminal prostate epithelial cell | urine cfRNA vs. plasma cfRNA | 3.56 E-05 |
| Proximal tubule | urine cfRNA vs. plasma cfRNA | 4.10 E-08 |

**Supplementary Table 3:** Internal standards added during metabolite extraction* prior to untargeted metabolomics analysis.

|  | Compound | Source | Product# |
| --- | --- | --- | --- |
| 1 | 1-Methylnicotinamide-d3 Iodine | TRC | M323237 |
| 2 | Carnitine-d3 (chloride) | Cayman Chemical | 26565 |
| 3 | Choline Chloride (Trimethyl-D9, 98%) | CAMBRIDGE ISOTOPE LABORATORIES, INC. | DLM-549-1 |
| 4 | Creatinine-d3 | Cayman Chemical | 16763 |
| 5 | CUDA* | Cayman Chemical | 10007923 |
| 6 | DL-Alanine-3,3,3-d3; 99.8% | CDN Isotopes | D-1462 |
| 7 | DL-Aspartic-2,3,3-d3 Acid 98% | CDN Isotopes | D-1953 |
| 8 | DL-Glutamic-2,4,4-d3 Acid 98.6% | CDN Isotopes | D-1196 |
| 9 | L-Acetylcarnitine-d3 (chloride) | Cayman Chemical | 26564 |
| 10 | L-Arginine-2,3,3,4,4,5,5-d7 HCl 98.1% | CDN Isotopes | D-7786 |
| 11 | L-Glutamine-(2,3,3,4,4-d5; 98.5%) | CDN Isotopes | D-2532 |
| 12 | L-Methionine (2,3,3,4,4-D5; Methyl-D3, 98%) | CAMBRIDGE ISOTOPE LABORATORIES, INC. | DLM-6797-0.1) |
| 13 | L-Phenylalanine(D8, 98%) | CAMBRIDGE ISOTOPE LABORATORIES, INC. | DLM-372-1 |
| 14 | Trimethylamine N-Oxide (D9, 98%) | CAMBRIDGE ISOTOPE LABORATORIES, INC. | DLM-4779-1 |
| 15 | Hippuric acid (Benzoyl- D5, 98%) | CAMBRIDGE ISOTOPE LABORATORIES, INC. | DLM-7703-01 |

\*denotes internal standard was added after extraction to control for LC-MS/MS injection efficiency

### Supplementary Note 1

We observed that the deconvolution error in the urine cfRNA samples was elevated and approached deconvolution error values observed while deconvolving GTEx tissues whose cell types were absent from TSP v1^1^. Inspection of the best-model gene expression predictions yielded poor prediction of *NEFH* and *KLK4*, genes that are highly expressed in luminal prostate epithelia relative to the other cell types in Tabula Sapiens, that were also poorly predicted by the support vector regression model and were therefore heavily penalized during RMSE calculation. The other top genes exhibiting high deconvolution error were shared between prostate cell types in Tabula Sapiens (Supplementary Fig. 5).

We bootstrapped 90% confidence intervals on the fractional estimation of cell type specific RNA (Methods) for these two cell types where we observed a positive interval in most male subjects (Supplementary Fig. 5c, d). We corroborated the presence of these cell types with signature scoring (Fig. 1d), indicating that the presence of this cell type was not a result of numerical artifact. We take this opportunity to emphasize that reporting outputted fractions as-is from a deconvolution program to assert relative cell type fractions can be limited without the ability to directly analyze the learned model, particularly when the final model coefficients were determined by a limited hyperparameter sweep and can additionally misassign transcriptionally similar cell type fractions from those that are insufficiently represented in the basis matrix column space.

We therefore implemented a fully open source, command line tool for individuals to deconvolve bulk RNA measurements - cellfracker (cell fraction estimator), which returns both the final regression model and the support vectors (genes) used to construct it, enabling direct analysis of the deconvolution results.

We further note that in two female donors, we observed large fractions of nominally prostatic cell types in their deconvolved urine transcriptomes and a 90% confidence interval spanning positive values. We additionally observed the largest deconvolution error across all samples for these two samples. These observations were congruent with the low, yet nonzero gene expression values we observed during signature scoring of the prostate epithelia (Fig. 1d).We therefore inspected the expression of these genes across the other cell types in Tabula Sapiens v1 basis matrix and in GTEx^1^ and the Human Protein Atlas RNA consensus dataset^2^; the latter two possess more comprehensive reference data for female reproductive tissues (e.g. ovary, cervix) than TSP v1. Although these genes exhibit very specific expression and high expression in the prostate relative to other tissues in the body, these genes also exhibit non-zero expression in female tissues and other select cell types. We therefore hypothesize that these genes play a role in other tissues that are not unique to men; though at-present they are frequently annotated as specific to the prostate in the literature.

Altogether, despite the limitations of deconvolving with a large column space owing to transcriptional similarity between cell types, we believe that unbiased deconvolution with the entirety of *Tabula Sapiens* (and subsequent model examination) can facilitate discovery of cell types contributing to a mixture that might otherwise be missed. However, care must be taken in interpretation given that *Tabula Sapiens* does not fully represent all cell types in the human body: for example, our observation of salivary gland cells most probably originates from secretory cells in the prostate responsible for producing seminal fluid. Indeed, without this approach, if we had restricted our basis matrix to urinary tract tissues, we would have missed the intestinal cells that we validated with known marker genes for these cell types.

